# Effects of dopamine D2 and opioid receptor antagonism on the trade-off between model-based and model-free behavior in healthy volunteers

**DOI:** 10.1101/2022.03.03.482871

**Authors:** Nace Mikus, Sebastian Korb, Claudia Massaccesi, Christian Gausterer, Irene Graf, Matthäus Willeit, Christoph Eisenegger, Claus Lamm, Giorgia Silani, Chris Mathys

## Abstract

Our daily behaviour requires a flexible arbitration between actions we do out of habit and actions that are directed towards a specific goal. Drugs that target opioid and dopamine receptors are notorious for inducing maladaptive habitual drug consumption, yet how the opioidergic and dopaminergic neurotransmitter systems contribute to the arbitration between habitual and goal-directed behaviour is poorly understood. By combining pharmacological challenges with a well-established decision-making task and a novel computational model, we show that the administration of the D2 dopamine receptor antagonist amisulpride led to an increase in goal-directed or ‘model-based’ relative to habitual or ‘model-free’ behaviour, whereas the non-selective opioid receptor antagonist naltrexone had no appreciable effect. These findings highlight the distinct functional contributions of dopamine and opioid receptors to goal-directed and habitual behaviour and support the notion that D2 receptor antagonists promote stabilisation of goal-relevant information.

## Introduction

Several theories of decision making postulate the existence of two distinct systems that drive our behaviour: a habitual system, which is automatic, reflexive and fast; and a goal-directed system, which is deliberative, reflective and effortful^1–3^. This dichotomy of systems has a computational analogue in ‘model-free’ and ‘model-based’ decision-making models^2,4^. A model-free agent simply selects actions that have led to rewarding outcomes in the past. This strategy is fast, computationally cheap, but can be inaccurate. A model-based agent uses an internal model of the environment to flexibly plan behavioural responses. This leads to more goal-oriented behaviour but is slower and relies on effortful cognitive control.

Studies investigating how individuals allocate control between the postulated two systems have shown that model-based control is increased when there is more to gain^5,6^, and decreased when cognitive resources are scarce^7,8^. Everyday behaviour can therefore be thought of as constant weighing of costs and benefits of applying model-based over model-free decision-making strategies^6,9,10^. Failure to exert cognitive control over habitual urges in order to avoid negative outcomes has been suggested to be a hallmark of substance addiction^2,11^. In support of this, studies show that decreased model-based control is linked to stimulant addiction^12^ and seems to constitute a transdiagnostic dimensional trait related to compulsive behaviour across clinical^12,13^ and non-clinical populations^14^. In light of increasing deaths from drug overdoses^15,16^, it is important to understand how different neurotransmitter systems affect the competition for cognitive resources when deciding between habitual and goal-directed actions.

Opiates, psychostimulants, and most other drugs of abuse increase the release of dopamine along the mesolimbic pathway^17,18^, a circuit that plays a central role in reinforcement learning^19^. On top of this, the reinforcing properties of addictive drugs also depend on their ability to activate the *µ*opioid receptors^20–22^. This invites the question whether the drift from goal-directed towards habitual control is enabled by dopamine and opioid receptors via a common neural pathway.

Yet, despite this question’s clinical relevance, causal investigations of the involvement of dopamine and opioid receptors in the trade-off between the two systems in humans are lacking. The evidence for the involvement of both dopamine and opioid receptors in forming and overcoming habits is restricted to preclinical studies on animals^18,22^ and to studies on patients with substance disorder^23,24^ with poor generalisability due to altered reward processing following chronic drug use^25^. Moreover, pharmacological studies in healthy volunteers are limited to unspecific dopamine receptor agonists and produced conflicting results^26,27^. The variability in these results may be due to dopamine’s involvement both in habit-formation as well as cognitive enhancement^28^, whereby its effect likely depends on the binding site^29^. Both D1 and D2 dopamine receptor families (called D1 and D2 in what follows) are involved in habit formation^30^ but appear to have opposing contributions to goal-stabilisation. D1 dopamine receptors in the prefrontal cortex enable maintenance of goal-relevant information and working memory, while D2 dopamine receptor activity disrupts prefrontal representations and supports cognitive flexibility^31–35^.

To address the limitations of previous studies, we directly compared the effects of the highly selective D2 dopamine receptor antagonist amisulpride with the opioid receptor antagonist naltrexone on the model-based/model-free trade-off in healthy volunteers. We hypothesized that both opioid and D2 antagonists would reduce habitual strategies and that blocking D2 receptors could also support model-based behaviour by stabilizing goal-relevant information^36^.

We tested 112 participants with a deterministic version of the two-step task^4,37^, at baseline and after administering amisulpride (N = 38, 400 mg), naltrexone (N = 39, 50 mg), or placebo (N = 35) in a randomized, double-blind, between-subject design (Fig. 1). The two-step task is a well-established incentivized paradigm, where participants need to make choices at the first stage, that influence their outcomes in the second stage. Each trial of the task starts in one of two possible first-stage states, each featuring a pair of spaceships (Fig. 2a). Upon choosing a spaceship, participants were taken to one of two planets in the second stage, where they encountered an alien that gave them points, which were converted to money at the end of the experiment. In each pair of spaceships, one spaceship flew to the red planet and one to the green. This deterministic mapping from spaceships to planets, as well as which spaceships were paired together, stayed constant throughout the task.

**Fig. 1.**
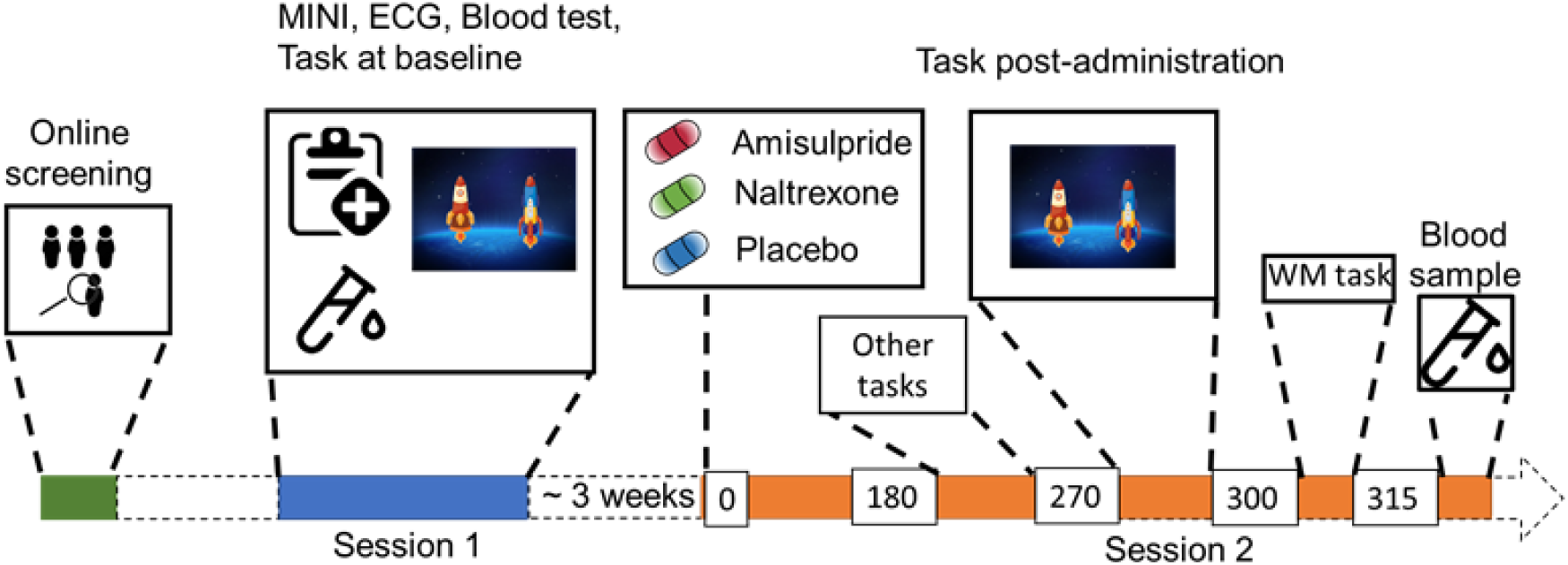
Study Procedure. After an initial online screening, participants were invited to the lab for a first visit (Session 1), where they were subjected to a medical screening, before playing the two-step task for the first time. If they fulfilled the study criteria, they were invited for another visit (Session 2), where they received either 400 mg of amisulpride, 50 mg of naltrexone, or placebo (mannitol). After 180 minutes of waiting time, participants started with the test battery. Approximately 270 minutes after drug intake, participants performed the two-step task the second time, followed by a Reading Span (working memory) task, and a blood draw to determine amisulpride serum levels.

**Fig. 2.**
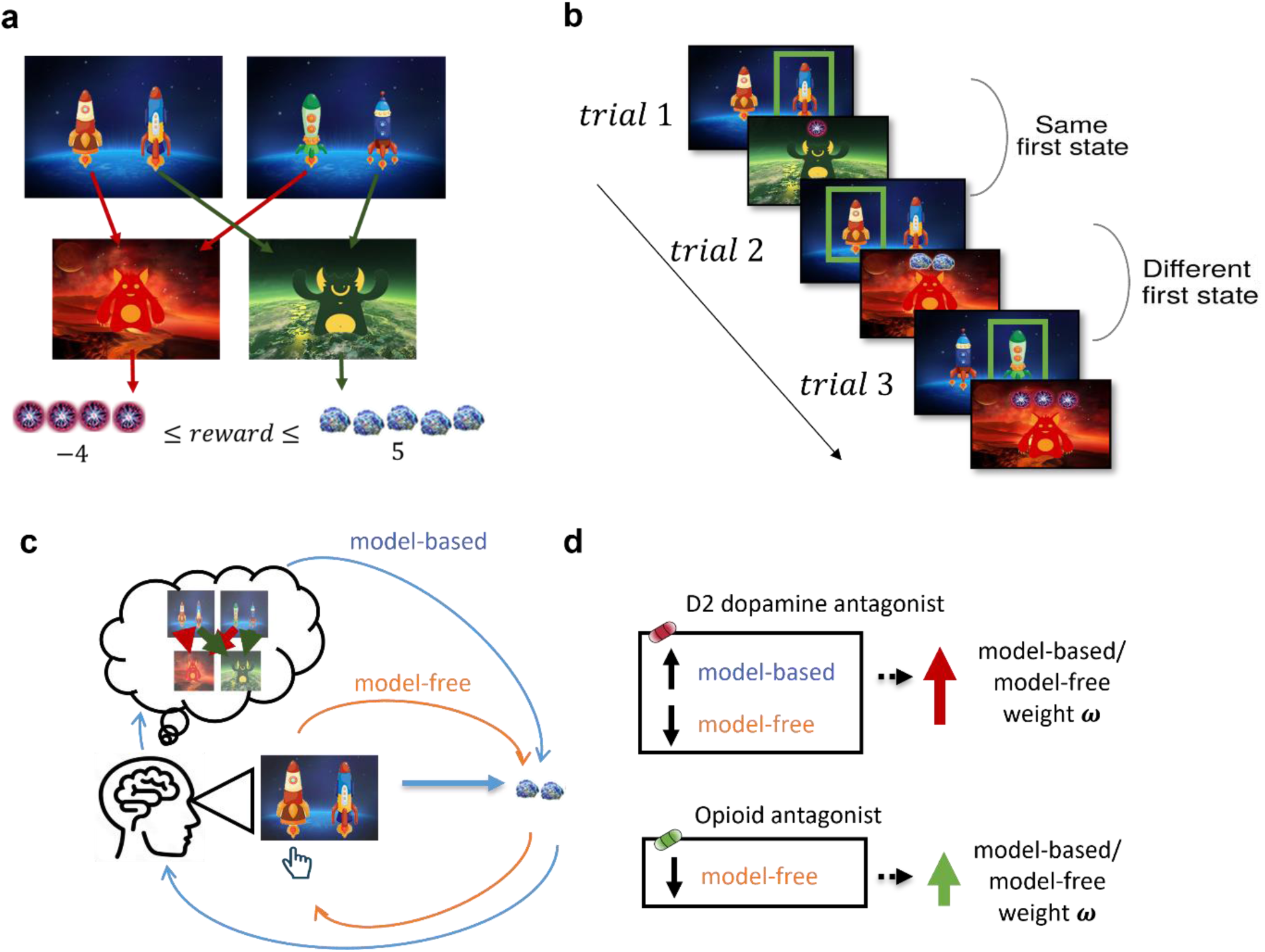
Task Design and Hypotheses. **a**, In the first stage of the trial, the participants were presented with one of the two pairs of spaceships. Each of the spaceships flew to one of two planets in the second stage, where they encountered an alien that gave them points. The transition from each of the four spaceships to the planets was deterministic and stayed the same throughout the experiment. The points each alien gave changed independently according to a discretized Gaussian random walk with bouncing boundaries at -4 and 5. **b**, Behaviour in the task showcasing trials where the previous first-stage state (spaceship pair) was the same (trial 2) and trials where it was different (trial 3). High number of points should encourage choosing the spaceship that flies to the same planet, what we term as “staying with the previous choice”. Note that in the featured example, the participant after receiving -1 point in the trial 1 opted against staying with the previous choice in the next trial, and after receiving 2 points in trial 2 opted for staying with the previous choice, by choosing the spaceship, that flew to the same planet. **c**, Model-free behaviour is defined by simple choosing the spaceship if it previously led to rewarding outcomes. In contrast, when behaving in a model-based manner, we choose the spaceship based on the planet it flies to, keeping the mapping from spaceships to planets in mind. **d**, We hypothesized that both amisulpride and naltrexone will increase the weight on model-based vs. model-free behaviour, however amisulpride would on top of that also support the model-based strategy.

However, the points received on each planet changed independently with a Gaussian random walk. When playing the game, participants could therefore simply repeat the choice of spaceship that previously led to positive outcomes, regardless of which planet it was associated with (model-free behaviour, Fig. 2b, c). Or they could attempt to remember which spaceship flew to which planet and choose the spaceship based on its associated planet (model-based behaviour, Fig. 2b, c). The task tried to mimic real-life decision-making dilemmas, where we can use our knowledge about the world (e.g., a potentially dangerous virus is circulating) to override a habitual action (e.g., shaking hands when greeting someone) with a more appropriate one (e.g., greeting with an elbow bump or a bow).

As an initial straightforward approach to dissociate model-based and model-free behaviour, we first examined how previous reward points affected the probability to stay with the previous choice. We then turned to computational modelling and explicitly modelled the relative weight, *ω*, of model-based compared to model-free behaviour and disentangled it from other processes that might affect participants’ choices, such as exploration and reward devaluation. Based on our hypotheses, we expected that both amisulpride and naltrexone would increase the weighting parameter *ω*, but that this difference would be more pronounced for amisulpride (Fig. 2d).

To assess any effects of the two pharmacological compounds on cognitive control, we also collected data on a working memory task. Finally, since a systemic administration of pharmacological compounds does not on its own allow for definitive conclusions about the specific neurobiological mechanisms involved in the decision-making process, we additionally explored how four genetic markers of baseline dopamine function moderate potential drug effects. The corresponding results will serve to direct future hypothesis-driven research. The data and the analysis scripts are available open-access at https://github.com/nacemikus/mbmf-da-op.git.

## Results

### Effects of dopaminergic and opioidergic antagonism on staying with previous choices

One way to quantify the trade-off between the two systems is by looking at the interaction effect of previous points on staying behaviour in trials where knowledge about the task structure is irrelevant (same first stage state as in the previous trial) with trials where it is relevant (different first stage states as in the previous trial). On average previous points increased the likelihood to stay with a previous choice (β_*logods*_= 0.402, 95% CI [0.251, 0.553], P(β_*logodds*_<0) < 10e-3, Fig. 3), however this was significantly less the case in trials with different vs. the same first stage states (β_*logods*_ = -0.204 (95% CI [-0.263, -0.146], P(β_*logodds*_<0) < 10e-3). This indicates that participants often failed to consider the mapping from spaceship to planets when making choices in trials where the first stage state differed from the previous trial.

**Fig. 3.**
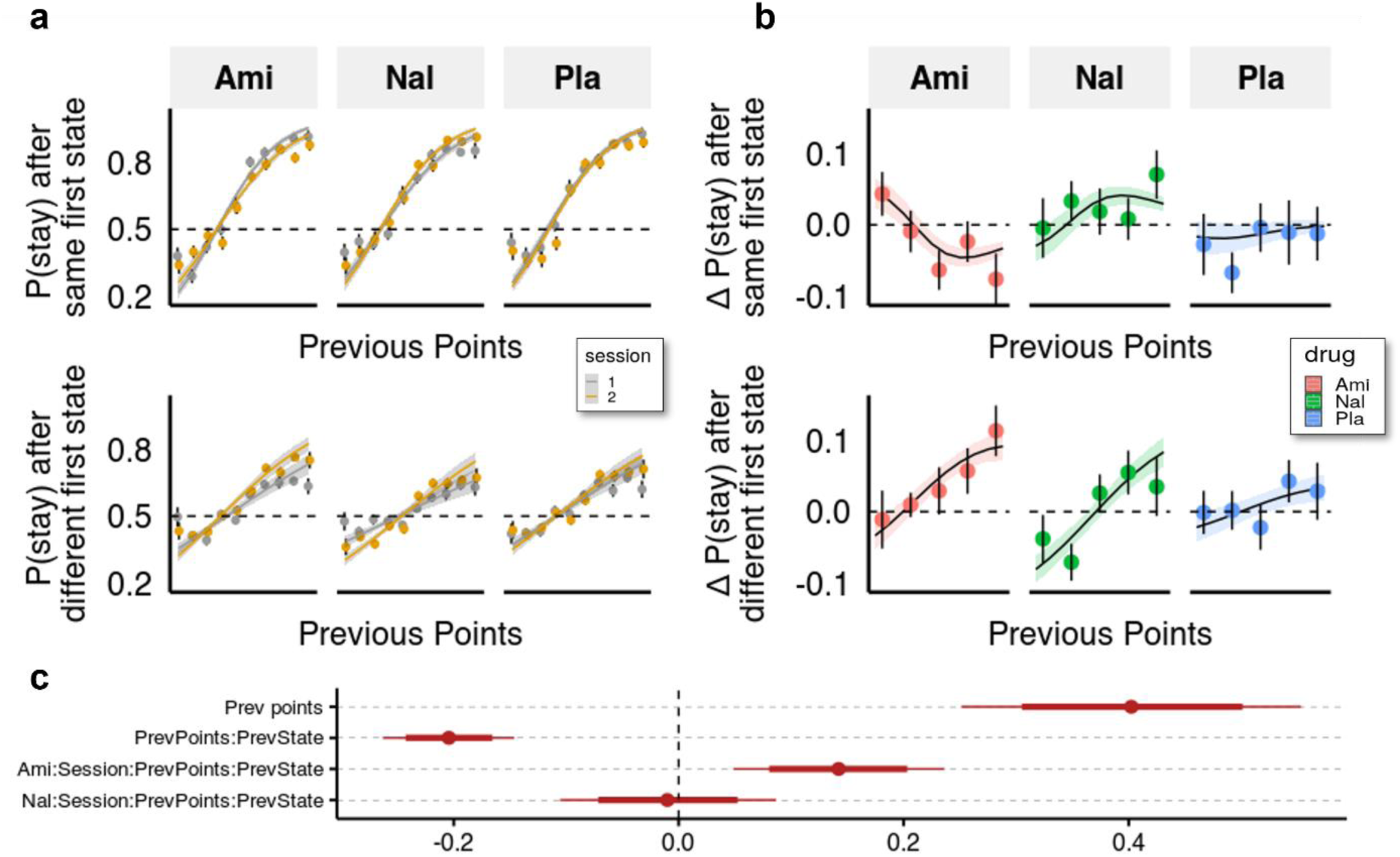
Behavioural analysis. **a**, We used a hierarchical Bayesian logistic regression model to analyse how the probability to stay with the previous choice depended on previous points, previous first-stage state, session, and drug administration and their interaction. We allowed the intercept and the slopes for previous first-stage state and session to vary by participant (full table of coefficients in Supplementary Table 1). Points and error bars depict mean proportions and standard errors in trials where the participants stayed with their previous choice for each previous point averaged within session and drug group. Lines and ribbons depict the mean estimates and 80% credible intervals (CI). When encountering a trial with the same first-stage state as in its preceding trial, participants were 1.495 times (95% CI [1.286, 1.739], P(β_*logodds*_ < 0) < 10e-3) more likely to stay with their previous choice for each additional previous point (β_*logodds*_ = 0.402, 95% CI [0.251, 0.553], P(β_*logodds*_ < 0) < 10e-3). In contrast, in trials where the first-stage state was different from the preceding trial, the odds (on the logarithmic scale) of repeating the previous choice of spaceship were reduced, with each additional point earned in the previous trial, by -0.204 (95% CI [-0.263, -0.146], P(β_*logodds*_ > 0) < 10e-3), and participants were only 1.219 times (95% CI [1.053, 1.410], P(β_*logodds*_ < 0) < 10e-3) more likely to stay with their choice for each additional previous point. This indicates that participants often failed to consider the mapping from spaceship to planets when making choices in trials where the first-stage state differed from the previous trial. The effects of the drugs can be seen by the different slopes of previous points in the two sessions depicted for both trial types. **b**, Differences in staying behaviour between sessions, binned into 5 different reward levels for clarity. Means with standard errors overlayed with means and 80% CI of estimated posterior distributions. **c**, Means with 80% and 95% CIs of effect sizes (in logodds space) of selected regression coefficients of the hierarchical logistic regression model predicting staying behaviour from previous points (PrevPoints), previous first-stage states (PrevState), session, drug administrations, and all the interactions between them. Ami, amisulpiride; Nal, naltrexone, Pla, placebo.

**Fig. 4.**
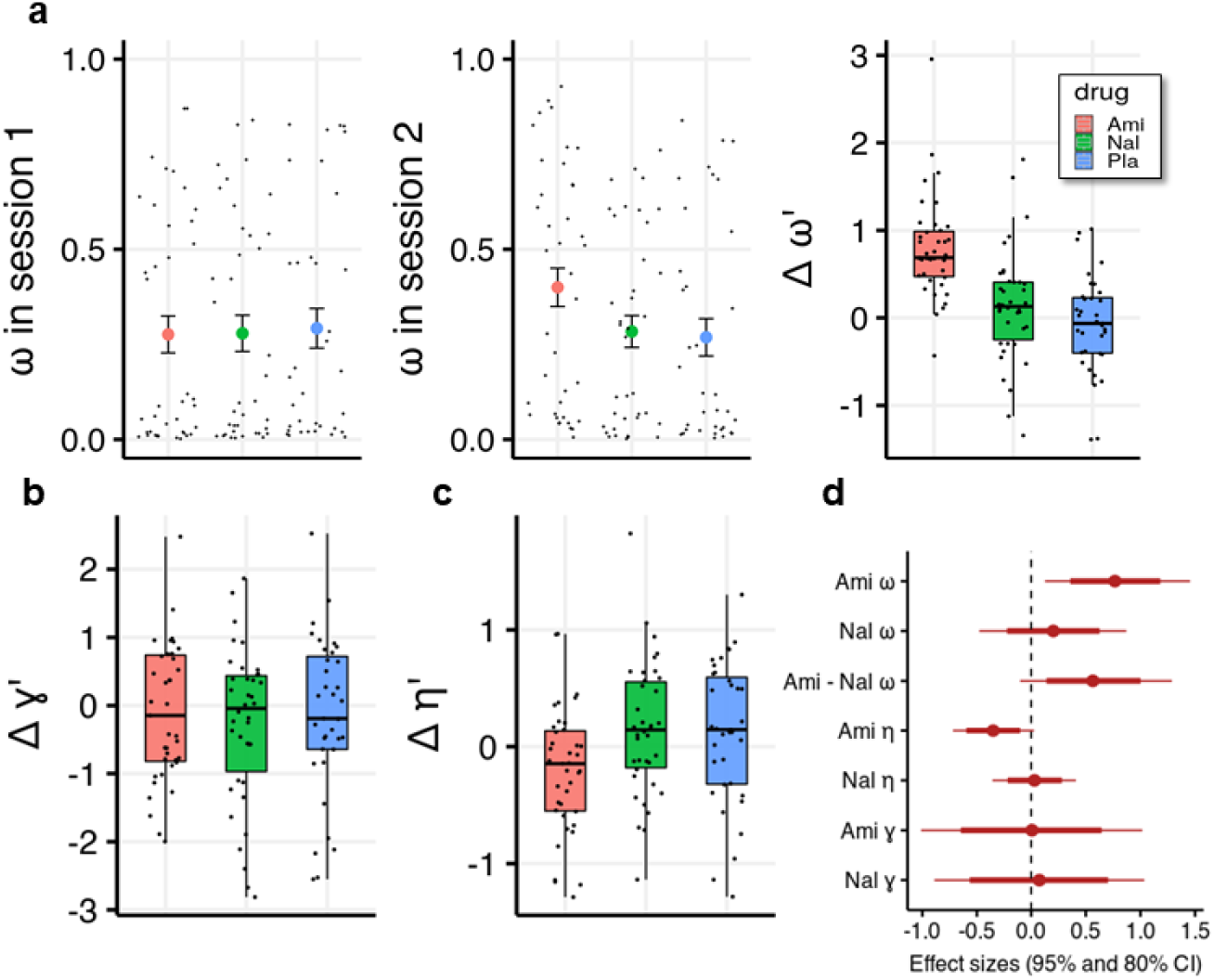
Effects of amisulpride and naltrexone on the model parameters. The best performing model (M1) is described with three free parameters, *ω* (the degree of model-based vs model-free value contributions to choice), *γ* (the degree of devaluation of unencountered spaceships), and *η* (the inverse temperature in the softmax mapping from values to probabilities). The parameters for both sessions and effects of drug treatments were estimated in one hierarchical model. **a**, Going from session 1 to session 2, amisulpride administration led to higher estimations of *ω*, and therefore increased model-based relative to model-free control. The difference between sessions of the model-based weights is shown in parameter estimation space (hence the prime). **b**, Difference in the parameter *γ*. **c**, Difference in the inverse temperature parameter. Lower values mean higher exploration. **d**, Posterior distributions of treatment effects on group-level mean session differences. Ami, amisulpiride; Nal, naltrexone, Pla, placebo.

Compared to placebo, amisulpride significantly increased the difference between the effects of previous points on staying behaviour in different vs. same first state trials, as indicated by a significant four-way interaction (β_*logodds* =_ 0.142, 95% CI [0.059, 0.234], P(β_*logodds*_<0) < 10e-3, Fig. 3). Interestingly, when looking at each type of trial separately, we found that amisulpride both decreased the effect of previous reward points on the probability to stay when the first-stage state was the same (β_*logodds* =_ -0.097, 95% CI = [-0.175 -0.018], P(β_*logodds*_>0) = 0.008), as well as increased it with a trend when the first-stage state was different (β_*logodds* =_ 0.046 (95% CI [-0.007, 0.098], P(β_*logodds*_<0) = 0.048). For naltrexone, the log odds estimate of the four-way interaction coefficient was centred around -0.011 (95% CI [-0.105, 0.087], P(β_*logodds*_<0) = 0.584), indicating no marked difference between trial types.

**Table 1:** Prior distributions for the behavioural analysis

|  |  |
| --- | --- |
| Standard Deviations | $\sigma \sim \text{HalfCauchy}(0, 2)$ |
| Regression Coefficients | $\beta \sim N(0, 3)$ |
| Intercept | $\beta_0 \sim \text{Student}(3, 0, 10)$ |
| Prior for the correlation matrix | $R \sim \text{LKJcorr}(2)$ |

### Estimation of drug effects with computational modelling

We designed a novel computational model that reflects the structure of the task and embedded the model parameter estimation within a hierarchical Bayesian inference framework^38^. We defined a model (M1) where, similarly to the computational models previously used with this task^6,37^, the trade-off between the goal-directed and habitual components is captured by the weighting parameter *ω*, which embodies the degree to which the choice of participants on each trial is influenced by model-based (*ω* = 1), or model-free (*ω* = 0) subjective values of each of the spaceships. In this model, the subjective values of both the model-free and the model-based model components are defined as the last observed outcome following the choice of that spaceship. The crucial difference between the two components is that, whereas the model-free agent learns only by experiencing direct outcomes of spaceship selection, the model-based component always considers the deterministic mapping from spaceships to planets, and thus learns the subjective values of planets. We also include an inverse temperature parameter *η* (lower *η* indicates more explorative behaviour) and a *discounting parameter γ* that marks the degree of devaluation of non-chosen (and non-encountered) spaceships for the model-free component in each trial. The model was validated through parameter recovery and posterior predictive checks^38^ and then compared to two other models commonly used in the field (see Methods and Supplementary Fig. 1).

#### Dopaminergic antagonism increases model-based relative to model free control

We first observed that the more model-based choices the participants made, the more money they earned (*r* = 0.65, 95% CI [0.53, 0.76]). This amounts to a validity check of the task, which was designed to make cognitive control pay off (literally)^37^. Importantly, we found that under amisulpride the difference in *ω* between the two sessions is higher than in the placebo group (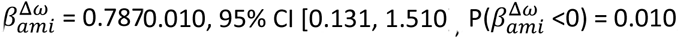, Fig. 3a), with effect size d = 0.758, (95% CrI [0.126, 1.455]). In contrast, there was no session difference in *ω* between naltrexone and placebo 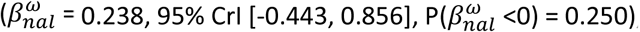, with a marginally significant difference with a moderate effect size between the effects of the two compounds 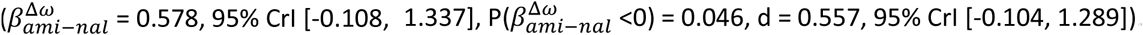.

#### Effects of drugs on other model parameters, mood and working memory

We found no evidence that either the blockade of dopamine D2 or opioid receptors influenced the devaluation parameter *γ* (Fig. 3b, d,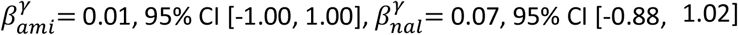). However, we found some evidence for amisulpride effects on the inverse temperature parameter *η* (Fig. 3c, d, 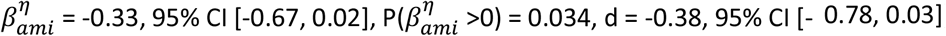). This would imply that amisulpride increases “explorative” choices or choices that the model would not predict based on estimated action values. To verify if our model performs worse for the amisulpride group we looked at how the pharmacological treatment predicts out-of-sample prediction accuracy and found no differences between groups (Supplementary Fig. 1f, Supplementary Table 2). Note also that the differences between sessions in *ω* and *η* are not negatively correlated (r = 0.12, 95% CI [-0.06, 0.30]), suggesting that the two effects are not related to each other.

To examine whether working memory capacity was affected by our drug manipulation, we analysed performance in the Reading Span task^39^ and found no evidence for drug effects on proportion of correct recalls 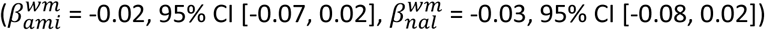. The increase in model-based behaviour could also be due to general effects on mood. We assessed participants’ mood on the day of drug administration twice, at drug intake and 3 hours later (Supplementary Table 3). A Bayesian linear model, predicting mood scores for positive and negative affect (PANAS)^40^, with varying intercepts for each participant, showed that in the placebo group there was a flattening of positive 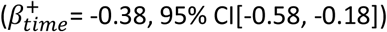 and, to a lesser degree, negative 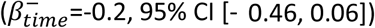 mood after 3 hours of testing (for the full table see Supplementary Table 4). The flattening in both affective domains was even more pronounced in the naltrexone group 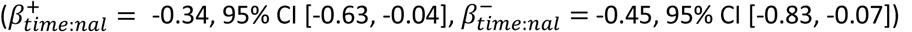, but was comparable to placebo in the amisulpride group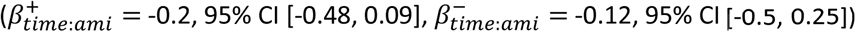. Finally, we ran a model predicting differences in *ω* between sessions, with working memory, mood and their interactions with the drug administration as predictors and found no effects (Supplementary Table 5), suggesting that the effects of amisulpride on *ω* were not due to effects of the drug on mood or working memory.

#### Explorative analysis of the interaction effects of pharmacological administration with genetically determined baseline dopamine function

In line with previous studies using similar paradigms, we used the *COMT* single nucleotide polymorphism (SNP) as a marker for prefrontal dopamine function, and *DARPP-32* SNP as a marker of striatal D1 receptor function^41–43^. We also investigated two genetic markers for striatal dopamine levels, namely the *Taq1a* SNP and the *DAT1* polymorphism, a 40 base-pair variable number tandem repeat polymorphism (VNTR) of the dopamine transporter^44–46^ (see Supplementary Note 2 for details on genotypes and how they relate to striatal and prefrontal dopamine function).

To estimate the moderating effects of the genotype variables, we ran again the hierarchical estimation of the parameters using shrinkage mixture priors (spike-and-slab) to regularize the parameter estimation^47^. The spike-and-slab prior pulls small effects towards zero and weakly regularizes non-zero effects^48,49^. To get a less conservative estimate of the effect, we ran a simple linear model predicting estimated changes of *ω* from one session to the next from drug and genotype interactions (Supplementary Fig. 2 a, b). We find that COMT moderated the effects of amisulpride (d = 0.597, 95% CrI [0.155, 1.049], P(d<0) = 0.005, Supplementary Fig. 2c), meaning that participants with genetic correlates of higher prefrontal dopamine were more likely to increase their model-based control following amisulpride administration. For naltrexone, we found a moderating effect of the Dat1 polymorphism (b = 1.077, 95% CrI [0.108, 2.038], P(d<0) = 0.015), and, to a lesser degree, of the other genotypes related to striatal dopamine function (Supplementary Fig. 2d). Note however that all the 95% posterior distributions of effect sizes from shrinkage priors contained zero (Supplementary Fig. 2b).

## Discussion

Drugs that stimulate opioid or dopamine neurotransmitter systems can lead to compulsive drug-taking, which can result either from increased reliance on habits or from failures to exert cognitive control over habitual urges. Here, we asked whether pharmacologically blocking opioid and dopamine receptors promotes the use of goal-directed over habitual control in healthy volunteers. Using the two-step task and a novel computational model, we found that blocking D2-like dopamine receptors increases the weight on model-based/model-free control, whereas blocking opioid receptors has no appreciable effect, with a marginally significant difference with a moderate effect size between the effects. The results from the computational model mirrored those obtained when analyzing the effects of previous points on staying with prior choices. These findings highlight the differential roles of the opioid and dopamine receptors in the model-based/model-free trade-off.

Prior research on dopamine’s involvement in the arbitration between the two systems has produced inconsistent results. For instance, elevated dopamine levels, either at baseline or following pharmacological treatments such as L-dopa, are related to increased model-based behaviour in some studies^26,50^, but show no correlation or even opposite effects in others^27,51^. This suggests that the effect of dopamine might depend on the relative contributions of the specific dopamine receptor subtypes or on the baseline differences in dopamine levels across different brain areas. It is well-established that dopamine enables cognitive control, mainly through prefrontal D1 receptors^32,52^. Striatal dopamine on the other hand is believed to control the gating of working memory content and therefore facilitates goal-relevant updating of information in the prefrontal cortex^35,53^. In the striatum, D2 receptor expressing cells are known to exert a tonic inhibitory effect on prefrontal and local circuits^54,55^. Higher D2 receptor activity disinhibits the cumulative basal ganglia output and promotes behavioural flexibility^35^. In line with this, D2 antagonists impair behaviour in tasks that require constant attentional set shifting^56^, but might improve performance in tasks where prefrontal goal representations are required^57^, such as the two step-task. For instance, D2 antagonism improves the ability of participants in certain cognitive tasks that require manipulation of task-relevant information in the working memory^36,53^, but does not improve working memory capacity, or memory retrieval^36,58,59^. This is supported by our data since we found no effect of amisulpride on working memory performance. Furthermore, an exploratory analysis of genetic data indicated that the effect of amisulpride might be moderated by prefrontal dopamine availability, which is known to be crucial in enabling model-based control^41,60^. This suggests that the D2 receptor antagonists’ effect on model-based behaviour is likely due to inhibiting disruptions of task-relevant prefrontal representations rather than to an increased capacity for cognitive control per se.

More recently, it has also been suggested that the striatal mediation of cognitive control reflects a willingness to exert cognitive effort, through a comparison of potential gains of applying cognitive control against the costs of doing so^29,61^. For example, a recent study showed that striatal dopamine synthesis capacity, measured with positron emission tomography, was related to higher motivation to exert cognitive effort^62^. Furthermore, in participants with low striatal presynaptic dopamine levels, sulpiride, an antipsychotic with a similar binding profile as amisulpride, increased the perceived benefits vs. costs of cognitive effort, an effect that was comparable to that of methylphenidate, a drug that increases dopamine availability in the synaptic cleft. D2 dopamine receptors are known to regulate dopamine reuptake, release, and synthesis through presynaptic auto-receptors^63^. Blocking D2 receptors could therefore boost the motivation to apply model-based control through increasing striatal dopamine levels. Interestingly, a recent study found that L-DOPA, a drug that increases presynaptic dopamine, did not increase model-based behaviour, but led to a general reduction of the effect of previous reward on choice, which was explained as reduced model-free behaviour, and higher noise.

In line with this, another explanation of amisulpride’s effects on the weighting parameter *ω* in our study is that they arise as compensation for disturbed model-free control. In previous studies, D2 antagonists led to poorer performance in simple reinforcement learning tasks^64,65^, reduced cue responding following Pavlovian conditioning, as well as preference for immediate rewards in a delayed discounting task^66^. In a task completed by the same cohort of participants prior to the two-step task, amisulpride reduced the willingness to exert physical effort to obtain immediate primary rewards^67^. In the present study, we found that amisulpride decreased the effect of previous points on staying behaviour in trials where the first-stage state was the same as in the previous trial, and therefore remembering the task structure was unnecessary. However, amisulpride increased the effect of previous points on choice in trials with different first stage states, suggesting that it did not disrupt learning overall. Another explanation for this is that amisulpride increased the capacity (or willingness) for model-based behaviour, but also increased choice exploration. Previous studies have implicated D2 receptors in exploration^46,68^ and we found some evidence of a slightly decreased inverse temperature parameter (i.e., more ‘noisy’ or explorative behaviour) in participants who received amisulpride. Furthermore, the drug effects on the exploration parameter and on the model-based/model-free weight were not correlated. Therefore, the increased exploration does not explain increased model-based/model-free weight and might reflect an orthogonal neurobiological effect.

Finally, participants could show reduced model-free learning that would reduce the effect or previous points on choice in trials with the same first-stage states, and at the same time reduce the misleading model-free action-outcome coupling that disrupts the model-based choice selection in trials where consecutive first states are different.

The lack of a pronounced effect of naltrexone on model-based/model-free behaviour is unlikely to be due to issues of dosage or timing, since 50 mg of naltrexone lead to a ∼90% of *µ* receptor occupancy even 48 hours post intake^69^, with *µ* receptors being the primary (but not only) opioid receptor type that the drug binds to. One possibility is that opioid receptors are not crucial for model-free learning required in this task. This is in opposition to previous studies showing that acute administration of naltrexone, comparably to amisulpride, causes a reduction of cue responsivity and reward impulsivity^66^, decreases effort to obtain immediate primary rewards^67^, and decreases the wanting of rewards^70^. Similarly, preclinical studies in rats show that naltrexone and naloxone, another opioid antagonist, decreased sucrose reinforced place preference^71,72^. Furthermore, both naltrexone and naloxone are used to reduce craving in patients with addiction^24^. However, there is also evidence that opioid receptors are important for goal-directed behaviour. In particular, a study in rats showed that opioid receptor blockade leads to decreased sensitivity to reward value and accelerated habitual control of actions^73^, and in a recent neurochemical imaging study in humans *µ* opioid receptor availability correlated with goal-directed behaviour in a loss-only version of the two-step task^51^. In fact, opioid agonists can in some cases lead to increased performance in cognitive control tasks^74^.

One explanation of our results is that naltrexone simply reduced the intrinsic value of reward and therefore decreased the motivation to exert cognitive effort. This would be in line with the observed flattening of both positive and negative affect after naltrexone administration compared to placebo. Finally, the lack of effects of naltrexone on the model-based/model-free trade-off might simply be too weak to detect in this study. Exploratory analysis of genetic data tentatively suggests that the effects might only be present in individuals with higher striatal dopamine levels, supporting the notion that habit formation through opioid receptors depends on striatal dopaminergic circuits.

An important limitation of the experimental approach used here is that it rests on the assumption that there are only two ways of learning, which moreover can be distinguished clearly. This assumption has been questioned in the reinforcement learning literature^75,76^, and mirrors the scepticism of the two-system division of decision making in cognitive psychology^77^. Although we made sure that participants understood what the task rules were and how they could maximize their gains, we cannot exclude the possibility that participants searched for alternative models to obtain rewards, such as different pressing patterns or simply favouring one stimulus over the other^75^. In fact, the slightly increased exploration parameter in the amisulpride group could be due to participants’ increased exploration of this model space.

In conclusion, we provide a first comparison of the contributions of the dopamine D2 and opioid receptors to arbitration between the model-based/model-free systems. Our results highlight the role of D2 dopamine antagonists in promoting goal-stability when alternative habitual choices are available. Furthermore, we show that opioid blockers in general do not promote model-based behaviour. These findings are a step forward in understanding how neuromodulators control the arbitration between habitual and goal-directed decision-making systems, which can on the long run be crucial for developing targeted pharmacological treatments for addiction and other disorders of compulsivity.

## Methods

### Procedure

The study took place in the Department of Psychiatry and Psychotherapy at the Medical University of Vienna. After initial online screening, the participants were invited for a first visit, where they underwent a physical examination (electrocardiogram, hemogram) and a psychiatric screening before being subjected to the two-step task that consisted of a 25 -minute automated training period followed by 200 trials of the task (see Figure 1 for the study outline). Participants eligible for the study were invited back to the clinic approximately 3 weeks after their first visit.

In their second visit participants received either placebo, 400 mg of amisulpride, or 50 mg of naltrexone. After a waiting period of 3 h, participants solved two additional tasks (explained elsewhere, see Korb et al., 2020) before performing the two-step task (on average 4 h 36 min, SD = 22 min, after the pill intake) as well as a working memory task. Waiting times and the doses of drugs were chosen based on previous pharmacological studies with the same compounds ^66^. To ensure comparable absorption, participants were asked to come with an empty stomach and received a standardized meal before pill intake. Participants filled out a questionnaire assessing mood and side effects right after pill intake and 3 hours later. There were no profound differences in side effects across the three drug groups, apart from a trend level effect of amisulpride on tiredeness (Supplementary Fig. 3). Blood plasma levels confirmed amisulpride levels above 212.6 *µ* g/l (mean (sd) = 548.8 (96.9)), in all participants of the amisulpride group.

### Participants

Data was collected from 120 volunteers and constituted a subset of volunteers involved in a larger study (Korb et al., 2020). All participants were assigned an initial screening code and a subject ID number. For six participants the task data from the first session were lost due to failures in assigning screening codes from the first session to subject ID numbers in the second. For two participants the task data from the second session were lost owing to technical issues, resulting in 112 participants included in the main analysis (see Supplementary Table 10 for exact subject numbers for each analysis). All participants had no history of drug abuse or other psychiatric disorders and were matched in age, sex, and BMI (Supplementary Table 8). The study was approved by the Ethical Committee of the Medical University of Vienna (EK N. 1393/2017) and in line with the Declaration of Helsinki ^78^. Participants received a monetary compensation of 90€ plus the extra money earned in the task.

### Genotypic analysis

Peripheral blood was collected by lancet and stored on Whatman FTA micro cards (Sigma-Aldrich). DNA was extracted using the QIAamp DNA Mini kit (Qiagen, Hilden, Germany). The VNTR polymorphism in the DAT1 gene was investigated by PCR with 5’-fluorescent-dye-labeled forward primer and automated detection of PCR products by capillary electrophoresis (details of the procedure provided in the supplement). The single base primer extension (SBE) method also known as SNaPshot minisequencing was applied for the typing of single nucleotide polymorphism (SNP) variants (details provided in supplement). Accordingly, five informative SNPs [ANKK1 (rs1800497), BDNF (rs6265), CDH13 (rs3784943), OPRM1 (rs1799971) and PPP1R1B (rs907094)] were analysed simultaneously applying a multiplex strategy for PCR and SNaPshot minisequencing of purified PCR products. Typing of Val158Met variants (rs4680) in the COMT gene was carried out separately, applying a singleplex approach for PCR and SNaPshot. The OPRM1 and BDNF SNPs were not included in the analysis in this task. Genetic data from three participants were lost. This led to the group distributions depicted in Supplementary Table 9. For details on the analysis see Supplementary Note 3).

### Task design

In the task, participants made an initial choice at stage one which took them to one of the two possible states (‘planets’) in stage two. There were two possible states in the first stage, each featuring a pair of spaceships. Each of the spaceships flew deterministically to one of the two planets where they encountered an alien who gave them either positive or negative points. The points in each planet changed according to a Gaussian random walk in the interval [-4, 5], rounded to whole numbers. Importantly, since both first-stage states led to the same two possible second-stage states, participants could transfer knowledge from the pair of spaceships in one first-stage state to the pair in the other. According to the conventional definition adopted here, a completely model-free agent relies entirely on its direct experience and will choose a spaceship based only on its reward history in previous trials featuring the same first-stage state, regardless of their experience with the other pair of spaceships. A completely model-based agent however will use the causal structure of the task to update the value of the spaceships based on which planet they fly to. This means that knowing that spaceship A and C fly to the same planet (but appear in different first step states) enables the model-based agent to learn about spaceship C by experiencing the outcome of its choice of spaceship A.

Participants had 2000 ms to choose in the first step and then again 2000 ms to press the space bar once they had encountered the alien. Each acquired point was translated to 4 Eurocents and added to the overall compensation of the participant at the end of the experiment. The story and a thorough training session were employed to increase the comprehension of the task, as done before^79^. To avoid all participants behaving in a model-based way^75^ we increased the difficulty of the task by dynamically changing the drift of the reward-determining Gaussian walks. The Gaussian random walks therefore had various drift rates (0.5, 1, and 2). We generated one trajectory and shuffled it around to create two different testing sessions of similar difficulty. Each participant first went through an automated rigorous explanation of the task followed by 25 practice trials before completing 200 trials of the task.

### Behavioural analysis

To regularize our inference, we used a hierarchical Bayesian approach in all our analyses. We report estimates in parameter estimation space and indicate the precision of our estimates with credible intervals^38,80^. We also report the proportion of the credible interval that is above or below zero. Model-agnostic analysis of behaviour focused on predicting the probability of staying with the previous choice (choose the spaceship that flies to the same planet as in the previous trial). Behaviour was analysed using the *brms* package in R^81^ which employs the probabilistic programming language Stan^82^. We fit a binomial model that predicted staying with the previous choice from reward obtained in the previous trial, modulated by the previous state (same or different), drug treatment, and session. The effects of session, previous state, and previous points were drawn from a multivariate normal distribution (were considered as correlated random effects). We report 95% credible intervals of estimates on the log-odds scale as well as in terms of odds (the ratio between probabilities of staying and switching) for a more comprehensive summary of the results. Parameters were estimated with 4 chains, each 3000 iterations (1000 warmup), with priors listed in Table 1.

The number of participants used in the analysis depend on whether both sessions were included and whether genetic data were used (see Supplementary Table 10 for an overview).

### Computational Models

We defined three computational models that define the subjective values *Q* of participants’ action *a*, in trial *t*, with a first stage state *s*, as the weighted average of model-based (*Q*_*MB*_) and model-free (*Q*_*MF*_) subjective values.

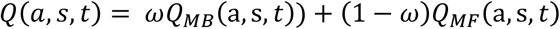

where *ω* is the weighting parameter (larger influence on choice of model-based values is indicated by *ω* being close to 1 and model-free control by *ω* being close to 0). The subjective values are then mapped on to probability of chosing *a*, and not *a*′, with the soft-max transformation:

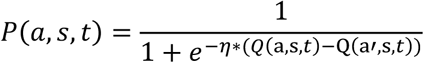

where *η* is the *inverse temperature* parameter, that determines the stochasticity of choices and the exploration-exploitation trade-off.

#### Deterministic learning model (M1)

Because the outcomes at the second stage are determined by Gaussian random walks, the objectively best prediction by the agent is the last encountered outcome. We defined a model (M1), where the model-free subjective value of each of the spaceships is the last encountered outcome following the choice of that spaceship, and the model-based subjective value of each spaceship is the subjective value of the last encountered outcome following the planet that this spaceship flies to. This means that the update equation of the model-free agent after receiving outcome *r*(*s*_2_, *t*) following an action *a*, in trial *t*, with first state *s*_1_, is defined as

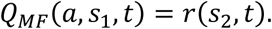

In the same trial, the value of the for the unchosen actions across both first level states are shrunk towards 0 with forgetting parameter *γ* ∈ [0,1]:

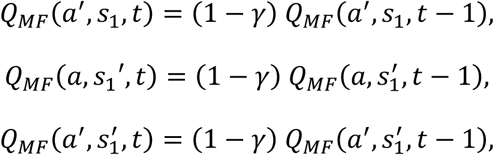

where *a*^′^ is the unchosen action and *s*_1_′ is the unencountered state. The model-based agent, on the other hand, after receiving outcome *r*(*t*) following an action *a*, in trial *t*, with first state *s*_1_, updates not only the experienced first stage state, but also the unencountered first stage which would have led to the same outcome:

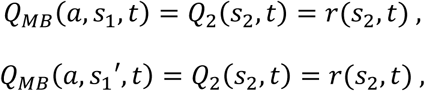

where, *Q*_2_(*s*_2_, *t*) is the subjective value of the second stage state that action *a* deterministically leads to.

#### *Dual-system reinforcement learning model*s (M2 and M3)

We compared our model M1 to two versions of a dual-system reinforcement learning model inspired by Kool et al (2016, 2017). In these models, the model-free agent learns the subjective values of spaceships and planets through a temporal difference-learning algorithm^9,83^. The model-free agent was defined by 3 free parameters: the learning rate at the first stage (α_1_), and the second stage (α_2_), where the eligibility trace (λ) determines the degree to which the outcome at the second stage retrospectively transfers to the first stage. In simple terms, the model increases (or decreases) the subjective value of an action at stage 1 proportionally to how positively (or negatively) surprising the outcome was, but discounted by the learning rate that describes the contributions of previous outcomes of that specific action. Conversely, the model-based agent is aware of the structure of the task. Specifically, in trial *t*, with first stage state *s*_1_ we define the model-free subjective value of action *a* as

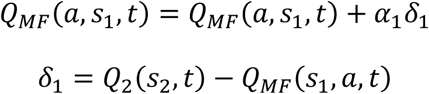

and the model-based subjective values of action *a* as

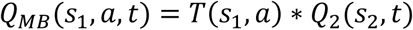

where *Q*_2_(*s*_2_, *t*) is the subjective value of the second stage *s*_2_ at time *t* and *T*(*s*_1_, *a*) is the transition matrix from first to second stage states. The agents learn the values of the planets, *Q*_2_, by

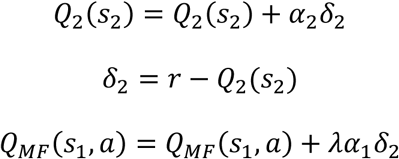

The learning rates at the two stages were either different (M2), or the same (M3).

### Model Estimation

We estimated the model parameters for both sessions in one hierarchical (multilevel) Bayesian model. This approach pools information across different levels (drug groups, and participants) and thus leads to more stable individual parameter estimates^84^, reduces overfitting^85^, and enables us to estimate in one model both individual and group level parameters as well as differences between sessions^86^. Models were implemented in Stan^82^ using R as the interface. Stan uses a Markov Chain Monte Carlo sampling method to describe posterior distributions of model parameters. We ran each candidate model with four independent chains and 3000 iterations (1000 warm-up). Convergence of sampling chains was estimated through the Gelman-Rubin 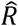 statistic^87^, whereby we considered 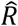 values smaller than 1.01 as acceptable.

For all subject level parameters (e.g., *ω*) we drew both the baseline (*ω*_0_) and the session difference (Δ*ω*), from a multivariate Gaussian prior. Specifically, for model M1:

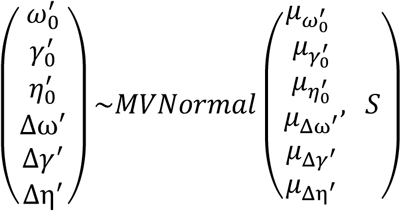

where *S* is the covariance matrix, that was factored into a diagonal matrix with standard deviations and the correlation matrix *R*^81,85^. The prime denotes the parameters in estimation space. The parameters that were fully constrained (i.e., *ω, α*_1_, *α*_2_, *λ, γ*) were estimated in the inverse probit space and the parameters that only had a lower bound (i.e., *η*) were estimated in log space. The hyper-priors for all group-level means were weakly informative, *µ*_*ω*_^′^ ∼ *N*(0,1), the prior for group-level standard deviations were σ_*ω*_^′^ ∼*HalfNormal*(0,1), and the prior for the correlation matrix was *R*∼*LKJcorr*(2). The parameters in the second session were therefore defined by their estimated baseline and difference between the sessions. For instance, the model-based weight *ω* in the second session for participant *i* was defined as:

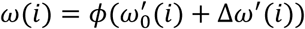

where *ϕ* is the cumulative distribution function of the standard normal distribution (inverse of probit). Effect sizes (d) are calculated by normalizing the relevant regression coefficients by the pooled standard deviation (square root of the sum of all relevant variance components^88,89^). In models where there are no random effects, this redcues to a Cohen’s d. In particular we defined the effect sizes by dividing the estimated difference between group means by the square root of the sum of the variance of both the baseline 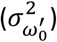 and the session difference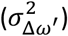. We used the same procedure for all three models.

### Model Comparison and Validation

We used the trial-based Leave-One-Out Information Criterion (LOOIC) to compare the three models using the loo package in R^90^. The LOOIC estimates out-of-sample predictive accuracy of each trial and is more informative than simpler point-estimate information criterions used commonly (such as the Akaike information criterion). Lower LOOIC scores indicate better prediction accuracy out of sample. Additionally, we compared models with bootstrapped pseudo Bayesian Model Average relative weights of the models that reflect the posterior probability of each model given the data ^91^. To validate the novel model M1, we first used the posterior means of the estimated parameters to simulate behaviour in both sessions. To see how well we can retrieve the model parameters we reran the parameter estimation on the simulated behaviour with the same model (Supplementary Fig. 1d). We also ran the same analysis of staying behaviour on this synthetic dataset and reproduced the behavioural plots from Fig. 3 (compare to Supplementary Fig. 1e). We used the same logistic hierarchical Bayesian model to statistically evaluate the crucial aspects of the behavioural analysis (Supplementary Fig. 1e, compare to Fig. 3c). To get the posterior predictive accuracy of the model we predicted the choice on each trial for each participant for 8000 samples drawn from the posterior distribution and then calculated the average accuracy for each participant.

### Estimation of the effects of the pharmacological treatment and genotypes

To statistically evaluate the effect of our treatment on all the parameters in the model, we included two regression terms when defining the group-level means of the difference between sessions:

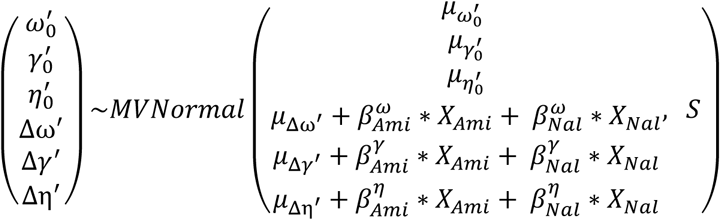

where 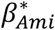 and 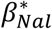 are coefficients drawn from prior distribution *N*(0,1.5) and *X*_*Ami*_ and *X*_*Nal*_ are dummy variables for the two drugs. To see which of the parameters are affected by the drug treatment we included group-level effects on all parameters.

To estimate the effects of the genotype on model-based behaviour, we again re-estimated the model adding the group level effects for each genotype to the session 1 estimates and interaction effects to the difference between session. All genotype variables were coded as binary and centred, except for the *COMT* which was coded as continuous and standardized. The hyper-mean of the model-based weights for the baseline was defined as:

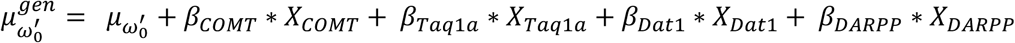

And for the session difference it was defined as:

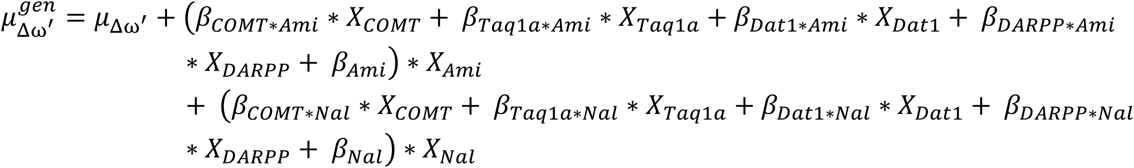

where *X*_*gene*_ are genotype variables and *β*_*gene*_ and *β*_*gene*drug*_ are coefficients drawn from a spike-and-slab prior, a prior that is a discrete mixture of a peaked prior around zero and a vague proper normal prior ^48,49^. Regression coefficients with weak effects are pulled towards zero, and the coefficients that are above zero are estimated with adaptive shrinkage. Specifically, we define the priors as

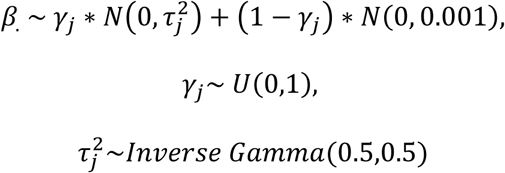

We only looked at the effect of genotypes on model-based weights. The effect sizes are calculated again by dividing the posterior distributions of regressors with the squared root of the sum of both the variance of the baseline *ω*_0_ as wall as the session difference. To get a less conservative estimate of the effects, we ran another model predicting estimated differences in *ω* across sessions from the drug variables and their interactions with the four genotype variables. Here, we standardized the response variable and used weakly informative priors (*N*(0,1)) on the regression coefficients and fat-tailed priors on the variance (*HalfCauchy*(0,2)). Effect sizes are calculated by dividing the posterior distributions of coefficients by the estimated standard deviation.

### Reading Span Task

We used an automated version of the Reading Span Task, where in each block participants saw 2 to 6 words serially presented that they had to recall by the end of the trial in any order. The words were interlaced with sentences that participants were instructed to judge as either making sense or not ^39^. Participants played 15 blocks. Correctly recalled items were calculated as a proportion within the block and then averaged across blocks. The effect of the drug treatment was calculated with a Bayesian linear model where the mean score was predicted by drug treatment.

## Acknowledgements

The study was supported by the Vienna Science and Technology Fund (WWTF) with a grant (CS15-003) awarded to Giorgia Silani and Christoph Eisenegger and a grant (VRG13-007) awarded to Christoph Eisenegger and Claus Lamm. We thank Prof. Boris Quednow for his advice on the study design. This work would not be possible were it not for the students who have carried out the data collection: Mani Erfanian Abdoust, Anne Franziska Braun, Raimund Bühler, Lena Drost, Manuel Czornik, Lisa Hollerith, Berit Hansen, Luise Huybrechts, Merit Pruin, Vera Ritter, Frederic Schwetz, Conrad Seewald, Carolin Waleew, Luca Wiltgen, Stephan Zillmer. We thank Catherine Hartley for sending us her version of the task implemented in matlab, and for allowing us to use the stimuli. We are grateful to Wouter Kool for making his version of the task implemented in JsPsych freely available on github. We also thank Lei Zhang and Lukas Lengersdorff for helpful discussions on the statistical procedures used in the manuscript.

## Notes

### Competing Interest Statement

The authors have declared no competing interest.

