## Supplementary Material for "Effects of dopamine D2 and opioid receptor antagonism on the trade-off between model-based and model-free behavior in healthy volunteers"

### Supplementary Figures

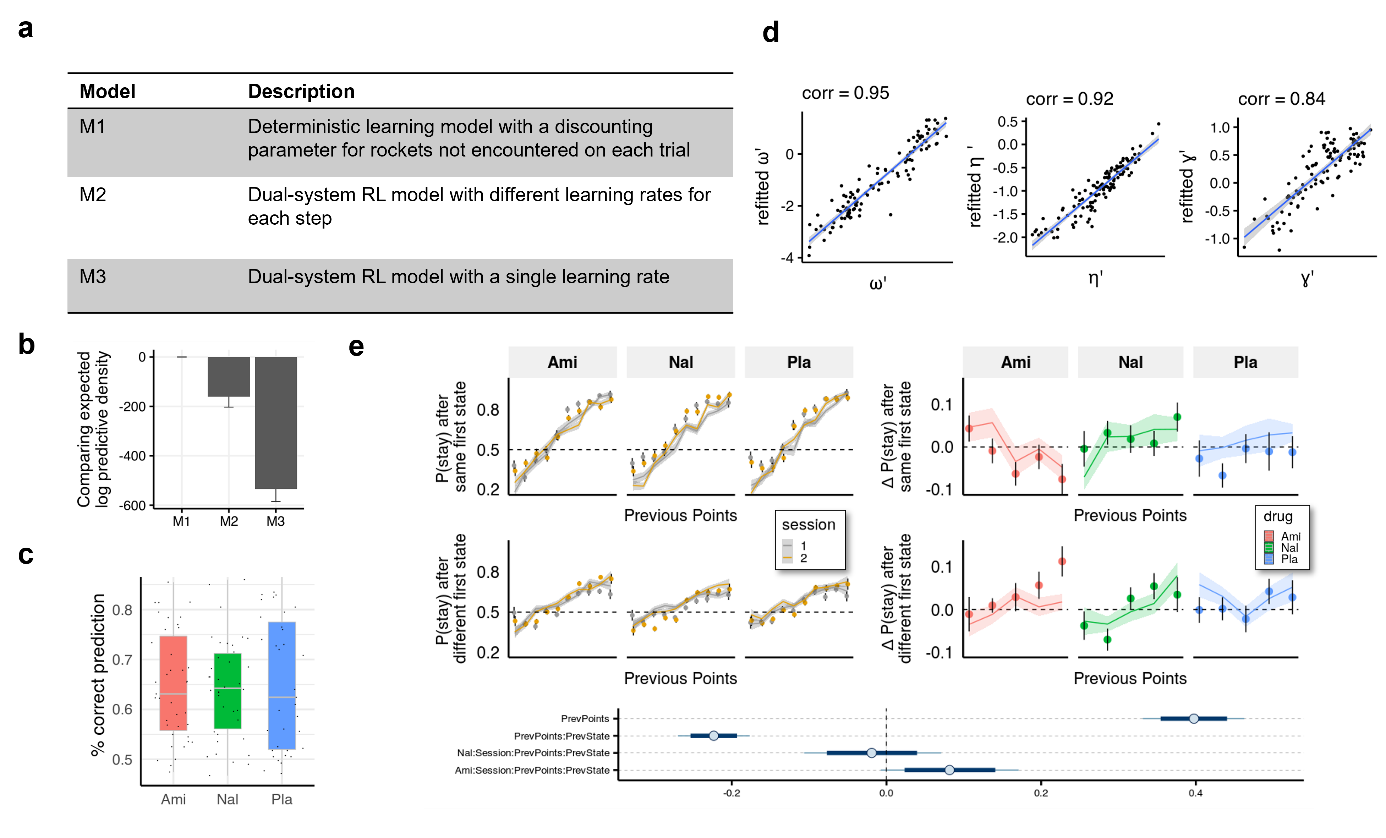

**Supplementary Fig. 1 | Computational modelling. a**, We compared our model to two dual-system reinforcement learning (RL) models used previously with this task^33,34^. In model M1 both model-free and model-based agents remember only the last outcome of a choice of spaceships, but only the model-based agent is aware of the relation between first-stage states. Models M2 and M3 are reinforcement learning-based models adapted from Kool et al (2016) with either separate (M2) or equal (M3) first and second stage learning rates. **b**, Models were compared using the leave-one-out information criterion (looic), where lower values indicate better out of sample trial-by-trial predictive performance. Refer to the Methods for weights from Bayesian Model Averaging of all models, indicating the posterior probability of each model given the data. Model M1 performs best in both metrics. **c**, For the winning model (M1) we calculated the posterior predictive accuracy averaged within participants and found that the mean (SD) posterior prediction accuracy was 65% (11%). **d,** Parameter recovery. **e**, Simulated data plotted to visually and statistically investigate whether the model captures the crucial aspects of behaviour and group differences (compare to Fig. 3).

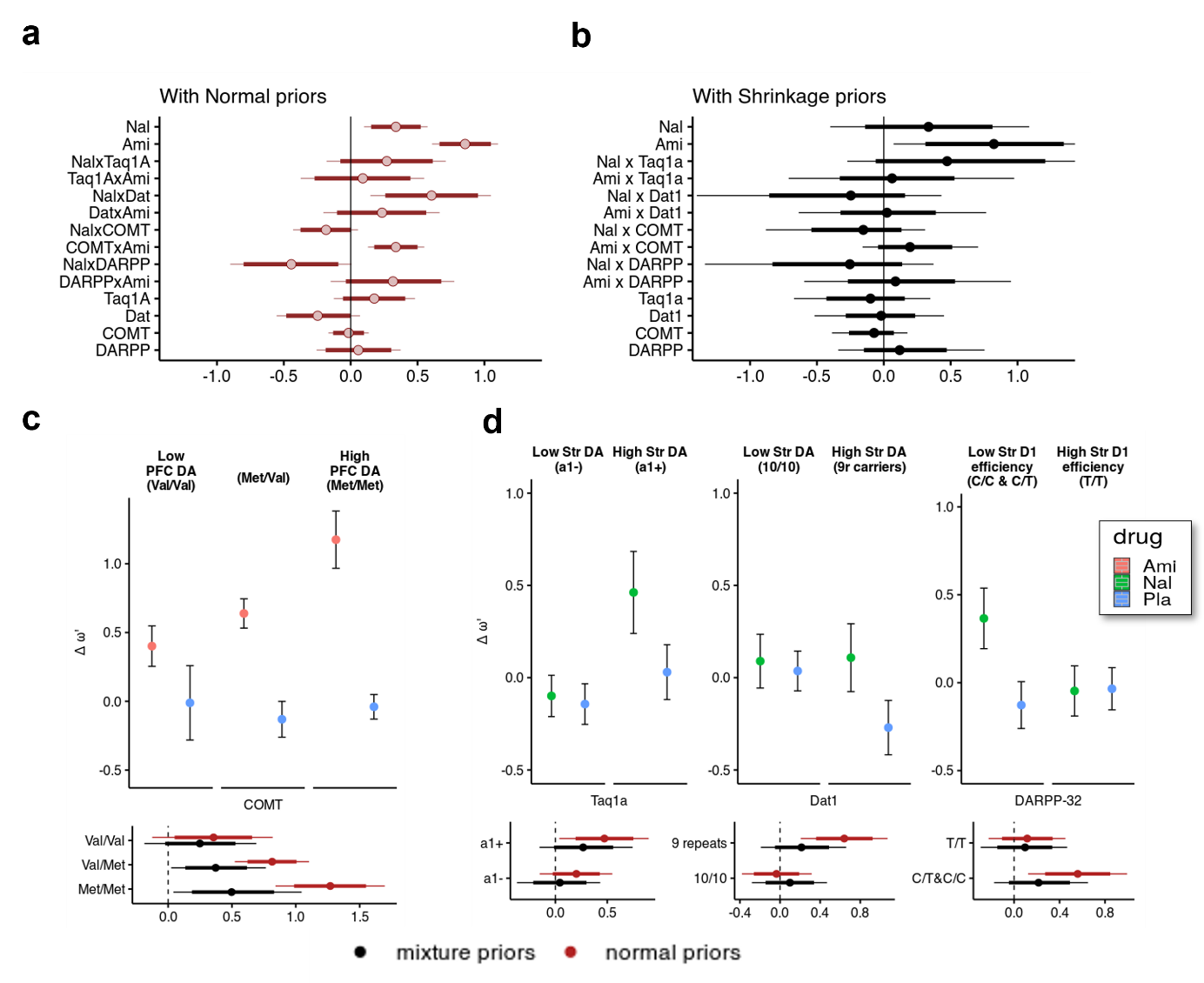

**Supplementary Fig. 2 | Interaction effects of genotypes and drug treatments on the weighting parameter** $\boldsymbol{\omega}$**. a, b** Posterior distributions of effect sizes (means, 95% and 80% intervals) estimating the genotypes x drug interactions using normal priors (a) and from the hierarchical model using mixture priors (b). **c**, The COMT polymorphism moderated amisulpride effects. We found that the more Met alleles individuals had (and putatively higher prefrontal dopamine availability), the higher the degree of model-based control (d = 0.335, 95% CI [0.09, 0.592], P(d<0) = 0.005, $d_{shrinkage}$ = 0.156, 95% CI [-0.159, 0.705], P($d_{shrinkage}$<0) = 0.19), with individuals with two Met alleles having the strongest effect of amisulpride on model-based behaviour (d = 1.270, 95% CI [0.841, 1.700], dshrinkage = 0.481, 95% CI [0.042, 1.050], P($d_{shrinkage}$<0) = 0.014). The moderating effects of striatal dopamine-related genotypes were less pronounced or minimal for the amisulpride group. **d**, Interaction effects of polymorphisms related to striatal function and naltrexone. We found that naltrexone increased $\omega$ in both A1 allele carriers (a1+) of the Taq1a polymorphism (d = 0.473, 95% CI [0.038, 0.899], P(d<0) = 0.016, d_shrinkage_ = 0.257, 95% CI [-0.153, 0.745], P(d_shrinkage_ <0.11)), as well as in participants with the 9-repeat polymorphism of the DAT1 polymorphism (d = 0.641, 95% CI [0.206, 1.070], P(d<0) = 0.002, d_shrinkage_ = 0.205, 95% CI [-0.192, 0.66], P(d_shrinkage_ <0) = 0.154). Furthermore, in participants with at least one C allele of the DARPP-32, naltrexone increased the weight on model-based behaviour (d = 0.561, 95% CI [0.123, 0.995], P(d<0) = 0.006, d_shrinkage_ =0.206, 95% CI [-0.176, 0.651], P(d_shrinkage_ <0) = 0.143). In other words, participants with putatively low D1 efficiency were more likely to employ model-based control following opioid antagonism. Participants with genotypes related to either higher dopamine availability or lower D1 receptor efficiency are more likely to have increased model-based behaviour following naltrexone administration. Ami, amisulpiride; Nal, naltrexone, Pla, placebo.

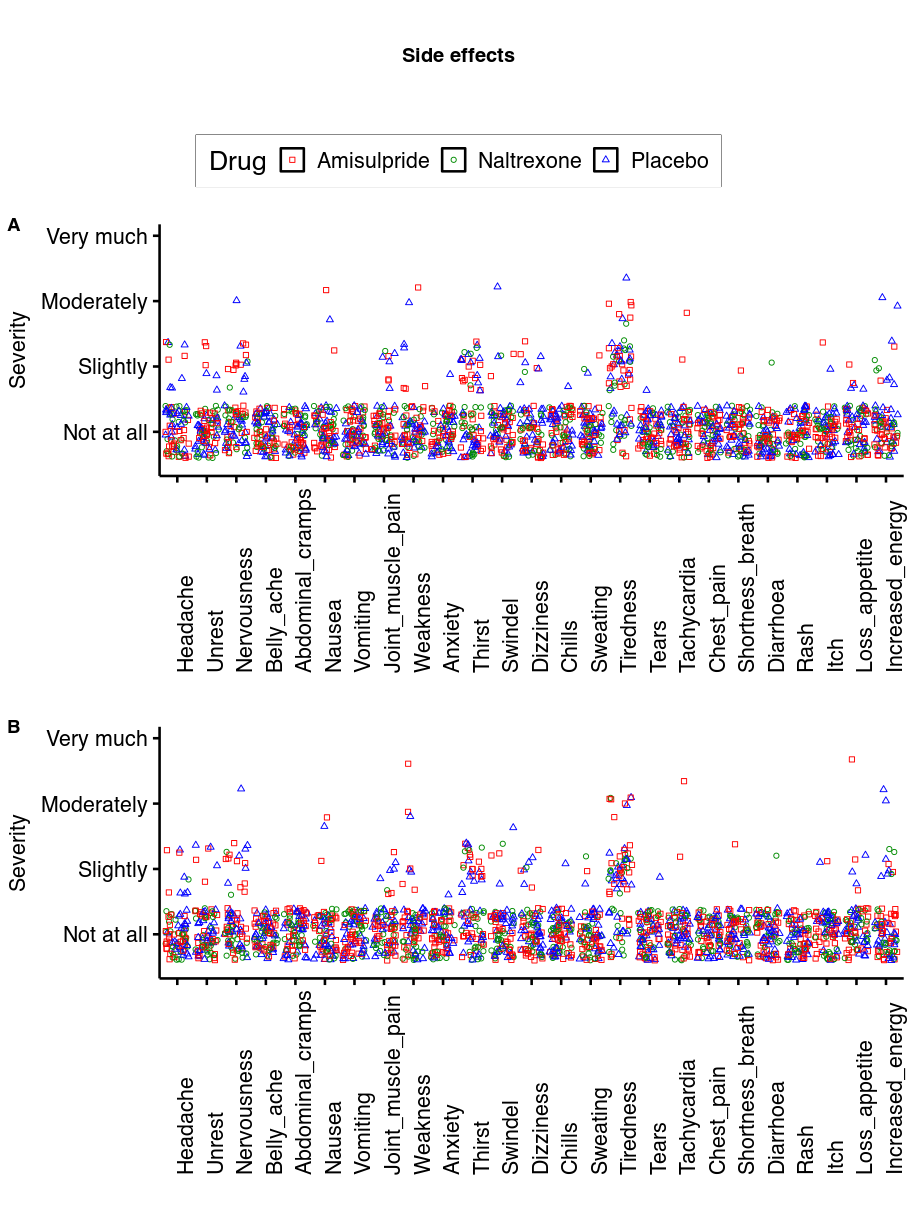

**Supplementary Fig. 3 | Side effects.** On most measures most participants rated the severity of the side effect as low as possible. The only measures where more than 50% of data points were above 1 was tiredness. Here, the effect of amisulpride was at trend level (b = 0.20, p = 0.06) and not significant for naltrexone (b = -0.15, p = 0.17).

### Supplementary Tables

| Stay ~ session * prev_points * prev_state_diff * Ami + session * prev_points * prev_state_diff * Nal +(session + prev_state_diff + prev_points \| ID) | Estimate | Q2.5 | Q97.5 |
| --- | --- | --- | --- |
| Intercept | 0.71 | 0.56 | 0.87 |
| session | -0.06 | -0.23 | 0.1 |
| prev_points | 0.44 | 0.35 | 0.53 |
| prev_state | -0.55 | -0.74 | -0.35 |
| Ami | -0.07 | -0.28 | 0.13 |
| Nal | -0.13 | -0.34 | 0.08 |
| session:prev_points | 0.01 | -0.05 | 0.06 |
| session:prev_state | 0.09 | -0.07 | 0.26 |
| prev_points:prev_state | -0.27 | -0.32 | -0.22 |
| session:Ami | -0.05 | -0.29 | 0.18 |
| prev_points:Ami | 0.04 | -0.09 | 0.17 |
| prev_state:Ami | 0.03 | -0.24 | 0.3 |
| session:Nal | 0.2 | -0.04 | 0.42 |
| prev_points:Nal | -0.06 | -0.2 | 0.06 |
| prev_state:Nal | 0 | -0.27 | 0.27 |
| session:prev_points:prev_state | 0.02 | -0.04 | 0.09 |
| session:prev_points:Ami | -0.1 | -0.18 | -0.02 |
| session:prev_state:Ami | 0.18 | -0.04 | 0.4 |
| prev_points:prev_state:Ami | -0.03 | -0.1 | 0.03 |
| session:prev_points:Nal | 0.06 | -0.02 | 0.14 |
| session:prev_state:Nal | -0.24 | -0.47 | -0.02 |
| prev_points:prev_state:Nal | 0.02 | -0.05 | 0.09 |
| session:prev_points:prev_state:Ami | 0.14 | 0.05 | 0.24 |
| session:prev_points:prev_state:Nal | -0.01 | -0.11 | 0.09 |

**Supplementary Table 1** | Estimates and CIs of fixed effects of the Bayesian logistic regression predicting staying behaviour. Q2.5 and Q97.5 are the 2.5% and 97.5% quantiles of the posterior parameter distribution. For details on how the posterior distributions were calculated refer to the code online (Ami = Amisulpride, Nal = Naltrexone).

| PosteriorPredictiveAccuracy ~ Ami + Nal | Estimate | Q2.5 | Q97.5 |
| --- | --- | --- | --- |
| Intercept | 0.69 | 0.51 | 0.87 |
| Ami | -0.05 | -0.3 | 0.2 |
| Nal | -0.07 | -0.32 | 0.17 |

**Supplementary Table 2 |**Results of a Bayesian logistic linear model predicting percentage correct from drug groups

|  | PANAS pos T1 | PANAS neg T1 | PANAS pos T2 | PANAS neg T2 |
| --- | --- | --- | --- | --- |
| Placebo | 29.5 (6.4) | 11.3 (1.6) | 27 (7.0) | 10.5 (0.9) |
| Amisulpride | 30.5 (5.6) | 12.2 (3.3) | 26.7 (6.2) | 10.9 (2.9) |
| Naltrexone | 30.1 (6.6) | 13.9 (7.0) | 25.4 (7.5) | 11.4 (4.4) |

**Supplementary Table 3** | Mood in mean and standard deviation at time of pill intake (T1) and 3 hours later (T2).

|  | PANAS Positive affect | | | PANAS Negative affect | | |
| --- | --- | --- | --- | --- | --- | --- |
| PANAS ~ time*ami + time*nal + (1\|ID) | Estimate | Q2.5 | Q97.5 | Estimate | Q2.5 | Q97.5 |
| Intercept | 0.19 | -0.11 | 0.49 | -0.1 | -0.4 | 0.2 |
| time | -0.38 | -0.58 | -0.18 | -0.2 | -0.46 | 0.06 |
| ami | 0.14 | -0.29 | 0.58 | 0.22 | -0.21 | 0.65 |
| nal | 0.05 | -0.38 | 0.48 | 0.65 | 0.22 | 1.09 |
| time:ami | -0.2 | -0.48 | 0.09 | -0.12 | -0.5 | 0.25 |
| time:nal | -0.34 | -0.63 | -0.04 | -0.45 | -0.83 | -0.07 |

**Supplementary Table** **4** | Results of a Bayesian linear analysis of both PANAS scales (centralized), with a varying intercept for each participant.

| $\Delta\omega$ ~ ($\Delta\mathrm{PANAS}_{\mathrm{neg}}$ + $\Delta PANAS_{pos}$ +BMI + WM + age) *(ami + nal) | **Estimate** | **Est.Error** | **Q2.5** | **Q97.5** |
| --- | --- | --- | --- | --- |
| **Intercept** | 0.059 | 0.154 | -0.234 | 0.362 |
| **nal_dummyNal** | 0.029 | 0.191 | -0.352 | 0.399 |
| **PANAS_negative_sess_diff_s** | -0.286 | 0.334 | -0.938 | 0.379 |
| **PANAS_positive_sess_diff_s** | -0.097 | 0.149 | -0.386 | 0.195 |
| **wm_s** | 0.125 | 0.133 | -0.143 | 0.382 |
| **Age_s** | -0.059 | 0.103 | -0.265 | 0.146 |
| **BMI _s** | 0.194 | 0.137 | -0.075 | 0.46 |
| **ami_dummyAmi** | **0.72** | **0.193** | **0.339** | **1.097** |
| **nal_dummyNal:PANAS_negative_sess_diff_s** | 0.3 | 0.376 | -0.431 | 1.046 |
| **nal_dummyNal:PANAS_positive_sess_diff_s** | -0.201 | 0.184 | -0.568 | 0.155 |
| **nal_dummyNal:wm_s** | -0.153 | 0.168 | -0.482 | 0.179 |
| **nal_dummyNal:Age_s** | -0.063 | 0.174 | -0.413 | 0.287 |
| **nal_dummyNal:BMI _s** | -0.229 | 0.177 | -0.575 | 0.119 |
| **PANAS_negative_sess_diff_s:ami_dummyAmi** | 0.57 | 0.42 | -0.244 | 1.411 |
| **PANAS_positive_sess_diff_s:ami_dummyAmi** | 0.136 | 0.185 | -0.238 | 0.498 |
| **wm_s:ami_dummyAmi** | -0.023 | 0.172 | -0.356 | 0.314 |
| **Age_s:ami_dummyAmi** | -0.104 | 0.159 | -0.425 | 0.204 |
| **BMI _s:ami_dummyAmi** | -0.067 | 0.184 | -0.425 | 0.29 |

**Supplementary Table** **5** | Results of a Bayesian linear analysis of predicting the difference in $\omega$ between sessions from working memory, age, BMI, and difference in mood for both PANAS scales (all centralized).

| **Drug** | **Genotype** | **level** | **d** | **Q2.5** | **Q97.5** | **P(d<0)** |
| --- | --- | --- | --- | --- | --- | --- |
| Ami | COMT | Met/Met | 0.481 | 0.042 | 1.046 | 0.014 |
|  |  | Val/Met | 0.366 | 0.024 | 0.766 | 0.017 |
|  |  | Val/Val | 0.244 | -0.185 | 0.691 | 0.12 |
|  | DARPP | C/T&C/C | 0.395 | 0.037 | 0.835 | 0.015 |
|  |  | T/T | 0.357 | -0.052 | 0.796 | 0.042 |
|  | DAT | 10-Oct | 0.383 | 0.017 | 0.798 | 0.02 |
|  |  | 9 repeats | 0.373 | -0.019 | 0.809 | 0.031 |
|  | Taq1a | a1- | 0.361 | 0.024 | 0.755 | 0.017 |
|  |  | a1+ | 0.387 | -0.033 | 0.897 | 0.033 |
| Nal | COMT | Met/Met | 0.078 | -0.413 | 0.499 | 0.364 |
|  |  | Val/Met | 0.165 | -0.18 | 0.518 | 0.173 |
|  |  | Val/Val | 0.237 | -0.225 | 0.875 | 0.156 |
|  | DARPP | C/T&C/C | 0.206 | -0.176 | 0.651 | 0.143 |
|  |  | T/T | 0.099 | -0.294 | 0.467 | 0.299 |
|  | DAT | 10-Oct | 0.1 | -0.281 | 0.468 | 0.293 |
|  |  | 9 repeats | 0.205 | -0.192 | 0.66 | 0.154 |
|  | Taq1a | a1- | 0.049 | -0.37 | 0.433 | 0.401 |
|  |  | a1+ | 0.257 | -0.153 | 0.745 | 0.113 |

**Supplementary Table 6 |** Results from the hierarchical model estimating the drug effects on model-based/model-free weight for each genotype group using shrinkage priors.

| $\Delta\omega\sim drug*\left( COMT+DARPP+DAT+Taq1A \right)$ | | | | | | |
| --- | --- | --- | --- | --- | --- | --- |
| **Drug** | **Genotype** | **level** | **d** | **Q2.5** | **Q97.5** | **P(d<0)** |
| Ami | COMT | Met/Met | 1.269 | 0.841 | 1.697 | <10e3 |
|  |  | Val/Met | 0.817 | 0.523 | 1.104 | <10e3 |
|  |  | Val/Val | 0.36 | -0.124 | 0.819 | 0.067 |
|  | DARPP | C/T&C/C | 0.698 | 0.253 | 1.131 | 0.001 |
|  |  | T/T | 1.014 | 0.654 | 1.373 | <10e3 |
|  | DAT | 10-Oct | 0.737 | 0.394 | 1.091 | <10e3 |
|  |  | 9 repeats | 0.971 | 0.544 | 1.401 | <10e3 |
|  | Taq1a | a1- | 0.808 | 0.47 | 1.15 | <10e3 |
|  |  | a1+ | 0.903 | 0.438 | 1.352 | <10e3 |
| Nal | COMT | Met/Met | 0.111 | -0.345 | 0.565 | 0.313 |
|  |  | Val/Met | 0.359 | 0.075 | 0.647 | 0.008 |
|  |  | Val/Val | 0.609 | 0.107 | 1.118 | 0.012 |
|  | DARPP | C/T&C/C | 0.561 | 0.123 | 0.995 | 0.006 |
|  |  | T/T | 0.118 | -0.225 | 0.452 | 0.247 |
|  | DAT | 10-Oct | -0.036 | -0.381 | 0.315 | 0.578 |
|  |  | 9 repeats | 0.641 | 0.206 | 1.072 | 0.002 |
|  | Taq1a | a1- | 0.204 | -0.15 | 0.549 | 0.127 |
|  |  | a1+ | 0.473 | 0.038 | 0.899 | 0.016 |

**Supplementary Table 7 |** Results from the linear model estimating the effects of drug and genotype interactions on difference of $\omega$ between sessions.

|  | N | BMI, m (sd) | Age, m (sd) | Sex(m,f) |
| --- | --- | --- | --- | --- |
| Placebo | 39 | 21.7 (2.2) | 23.2 (3.6) | 12,27 |
| Amisulpride | 39 | 22.7 (2.6) | 22.8 (2.8) | 14,26 |
| Naltrexone | 40 | 23.0 (2.4) | 23.2 (4.0) | 13,26 |

**Supplementary Table 8 |**Description of participants in terms of body mass index (BMI), age and sex with mean (m) and standard deviation (sd).

|  | COMT  Val/Val, Val/Met, Met/Met | DAT1  10/10, other | ANKK,  a1-, a1+ | DARPP  C/C+C/T, T/T |
| --- | --- | --- | --- | --- |
| Placebo | 9,14,12 | 23,12 | 20,15 | 13,22 |
| Amisulpride | 10,15,12 | 20,17 | 27,10 | 13,24 |
| Naltrexone | 6,20,11 | 20,17 | 24,13 | 13,24 |

**Supplementary Table 9** | Distribution of genotypes

|  | Behavioural Analysis with both sessions | Computational modelling with both sessions | Computational modelling, second session only | Computational modelling with both sessions and genetic data |
| --- | --- | --- | --- | --- |
| Placebo | 35 | 35 | 39 | 35 |
| Amisulpride | 38 | 38 | 39 | 37 |
| Naltrexone | 39 | 39 | 40 | 37 |

**Supplementary Table 10 |** Number of participants per drug group used in analysis

### Supplementary *Notes*

**Supplementary Note 1**

One of the peculiarities of our design is a baseline measure, and the fact that the participants were all introduced to the task under no administration thus avoiding potential effects of the treatment on task training. However, due to this design, we cannot completely exclude order effects. Although unlikely, it is possible that the effects of the treatment that we observe come indirectly from the effects of the two drugs on either skill transfer from the previous session, or simply on the effect of the drugs on the part of the experiment that preceded the task, which could mean, for instance, that participants under amisulpride were less tired from other tasks and therefore more willing to exert effort in the task presented here. Speaking against a general effect is the observation that we found no differences in mood between amisulpride and placebo.

**Supplementary Note 2: Description of genotypes and how they relate to baseline dopamine function**

*Taq1a* is a D2 dopamine receptor polymorphism with two allelic variants, A1 and A2. The A1 allele carriers (a1+) have reportedly higher dopamine synthesis rate^1^ that could potentially result in lower density of D2 receptors in the striatum^2,3^. The *Dat1* VNTR is a polymorphism of the dopamine transporter (DAT) that is involved in reuptaking dopamine from the synaptic cleft back into the cell and thus a regulator of tonic dopamine levels and occurs with highest density in the striatum^4^. The 9-repeat variant of the polymorphisms is associated with lower levels of DAT, and thus potentially higher extra-cellular dopamine levels^5,6^. Based on the above both an effect on baseline tendency for model-based behaviour as well as an augmentation of the drug effects could be expected in the group carrying at least one A1 allele (a1+ group) and at least one the 9-repeat variant of the DAT1 polymorphism (9 repeats group). As in previous studies, we used the valine-to-methionine (Val/Met) substitution in the Val^158^Met polymorphism of the *COMT* gene as a marker for prefrontal dopamine function^7,8^. COMT (or Catechol-O-methyl transferase) is an enzyme that has an effect on dopamine extracellular levels through metabolizing dopamine especially in cortical areas of the brain^9^. Participants with Met mutations have lower enzymatic activity of the *COMT* and have been associated with higher dopamine levels the prefrontal cortex^10^. And lastly, we used the *DARPP-32* as a marker of striatal D1 receptor function. T homozygotes (T/T allele carriers) have putatively higher efficiency of striatal D1 receptors^11^ and have been related to model-free learning ^12,13^.

To estimate the moderating effects of the genotype variables, we ran again the hierarchical estimation of the parameters, including group level variables for four genetic variables, as well as their interactions with the two drug treatments as modulators of the variable that estimates the difference in $\omega$ between the two sessions. We used shrinkage mixture priors to regularize the parameter estimation^54^. The spike-and-slab prior pulls small effects towards zero and regularizes non-zero effects^55,56^. With this analysis, there were no moderating effects of genotypes (with 95% CrIs not including zero, Supplementary Fig. 2 b). To get a less conservative estimate of the effect, we ran a simple linear model predicting estimated changes of $\omega$ from one session to the next from drug and genotype interactions (Supplementary Fig. 2 a). We found some evidence that COMT moderated the effects of amisulpride(d = 0.597, 95% CrI [0.155, 1.049], P(d<0) = 0.005, Supplementary Fig. 2 c), meaning that subjects with genetic correlates of higher prefrontal dopamine were more likely to increase their model-based control following amisulpride administration. For naltrexone, we found a moderating effect of the Dat1 polymorphism (b = 1.077, 95% CrI [0.108, 2.038], P(d<0) = 0.015) and of the DARPP ("b = -0.787, 95% CrI [-1.745, 0.173], P(d>0) = 0.053), but no support for the notion the prefrontal dopamine affects naltrexone’s effects on model-based/model-free trade-off (Supplementary Fig. 2 d). This would suggest that high striatal dopamine levels and low striatal D1 receptor efficiency enabled an increase in model-based behaviour after opioid receptor antagonism. Note that this analysis likely underestimates the true variance, but the results might nevertheless inform future hypothesis driven investigations.

**Supplementary Note 4: Details of Genotyping**

**I. Typing of the variable number tandem repeat (VNTR) polymorphism in the DAT1 gene**

Peripheral blood was collected by lancet and stored on Whatman FTA micro cards (Sigma-Aldrich). DNA was extracted using the QIAamp DNA Mini kit (Qiagen, Hilden, Germany) and eluted in a final volume of 50 μL of buffer AE (Qiagen, Hilden, Germany). Human DNA concentration was determined with the Quantifiler HP Quantification Kit (AB) on the Applied Biosystems (AB) 7500 real-time PCR instrument (Thermo Fisher Scientific, Waltham, MA). Template DNA (10 ng per sample) was subjected to PCR in a total reaction volume of 25 µL consisting of 1 × GeneAmp PCR Buffer (AB), 0.25 mM each dNTP, 2.5 U AmpliTaq Gold Polymerase (AB) and target specific primers (details are provided in Table S1). The following thermal protocol was applied using a Veriti 96-Well Thermal Cycler (AB): 35 amplification cycles at 95 °C for 30 seconds, 55 °C for 1 minute, and 72 °C for 1 minute. Before the first cycle, an initial “hot start” denaturation (5 minutes at 95°C) was included, and the last cycle was followed by a final extension step at 72 °C for 45 minutes. Aliquots of PCR products were diluted with Hi-Di formamide (AB) mixed with internal lane standard LIZ 600 v.2 (AB) and separated on the ABI 3500 Genetic Analyzer applying standard conditions. The number of repeats predicted by the GeneMapper ID-X software (AB) was in full agreement to the number of repeats determined by direct sequencing of PCR products using the BigDye Terminator Sequencing Kit v3.1 (AB) in selected DNA samples.

**Table S1. Primer set used for the typing of DAT1 VNTR repeat length polymorphisms**

| Marker | Location^a^ | Primer sequence 5’-3’ ^b^ | Dye ^c^ | Orientation | Conc.(nM)^d^ |
| --- | --- | --- | --- | --- | --- |
| DAT1 VNTR | chr5:1393559-1394008(-) | TGTGGTGTAGGGAACGGCCTGAGA | 6-FAM | forward | 400 |
|  |  | TGTTGGTCTGCAGGCTGCCTGCAT |  | reverse |  |

^a^ Chromosome number and genomic location of targeted sequence (orientation provided in brackets) according to UCSC version hg38 (<http://genome.ucsc.edu/>) is provided.

^b^ The non-specific primer tail is underlined in Italics.

^c^ 5’ Fluorescein (6-FAM)-labeled forward primer was used.

^d^ The final primer concentrations in the reaction mix is shown.

**II. Typing of single nucleotide polymorphisms (SNPs) by SNaPshot minisequencing**

Five informative SNPs [ANKK1 (rs1800497), BDNF (rs6265), CDH13 (rs3784943), OPRM1 (rs1799971) and PPP1R1B (rs907094)] were typed simultaneously applying a multiplex strategy for PCR and SNaPshot minisequencing of purified PCR products. Typing of Val158Met variants (rs4680) in the COMT gene was carried out separately, applying a singleplex approach for PCR and SNaPshot.

**Step 1: PCR and purification**

**1.1 Singleplex PCR (COMT)**

First, a 177 bp genomic fragment of the COMT gene harbouring the causative single nucleotide polymorphism (SNP rs4680) in its centre was targeted by PCR. The reaction mix contained 5 ng template DNA, 1 × GeneAmp PCR buffer (AB), 0.25 mM each dNTP, 2.5 units AmpliTaq Gold polymerase (AB) and specific primers (details provided in Table 2) in a total reaction volume of 25 µL. Thermal cycling was conducted using the Veriti cycler (AB) and the following conditions: 95 °C for 5 min; 35 cycles of 95 °C for 15 seconds, 59 °C for 30 seconds and 72 °C for 1 minute; final extension at 72 °C for 5 minutes. Excess of primers and unincorporated dNTPs were removed by adding 2 µL of ExoSAP-IT (Thermo Fisher Scientific) to each 5 µL PCR product. Reactions were incubated at 37˚C for 15 min followed by 80˚C for 15 min for enzyme deactivation.

**1.2 Multiplex PCR (ANKK1, BDNF, CDH13, OPRM1, PPP1R1B)**

Amplification by multiplex PCR was carried out in a total volume of 20 µL, containing 1x Phire Hot Start II PCR Master Mix (Thermo Fisher Scientific), PCR primers concentrations as specified in table 2 and template DNA (5 ng per sample). Thermal cycling was carried out on the Veriti thermal cycler (AB), and the following conditions: 98˚C pre-incubation step for 30 sec; 40 cycles of 98˚C denaturation for 10 sec, annealing at 62˚C for 30 sec and extension at 72˚C during 30 sec; followed by 1 min for final extension at 72˚C. PCR products were purified by ExoSAP-IT treatment.

**Table 2. Panel of loci and primer sets used for PCR**

| Marker^a^ | Location^b^ | Primer sequence 5’-3’ | Orientation | Conc.(nM)^c^ |
| --- | --- | --- | --- | --- |
| ANKK1 | chr11:113400010-113400194(-) | GACATGATGCCCTGCTTTCG | forward | 500 |
|  |  | CATCACGCAAATGTCCACGC | reverse |  |
| BDNF | chr11:27658331-27658494(-) | CAAACATCCGAGGACAAGGT | forward | 250 |
|  |  | CCGAACTTTCTGGTCCTCAT | reverse |  |
| CDH13 | chr16:83678696-83678883(+) | CAGTGCTGTGTTCCCAAATG | forward | 500 |
|  |  | CTGCGAATCCACAATACCTGT | reverse |  |
| COMT^a^ | chr22:19963623-19963799(+) | GGGCCTACTGTGGCTACTCA | forward | 400 |
|  |  | GCCCTTTTTCCAGGTCTGA | reverse |  |
| OPRM1 | chr6:154039591+154039751(+) | CCTTGGCGTACTCAAGTTGC | forward | 500 |
|  |  | CGTGATCATGGAGGGACTG | reverse |  |
| PPP1R1B | chr17:39634064-39634252(-) | CTCAGGTAGGGCTGAGTTCG | forward | 500 |
|  |  | CCTGAAGGTCATCAGGCAGT | reverse |  |

^a^ COMT primers only used in singleplex PCR; all other primers combined in a multiplex PCR.

^b^ Chromosome number and genomic location of targeted sequence (orientation provided in brackets)

according to UCSC version hg38 (http://genome.ucsc.edu/).

^c^ The final primer concentrations in the reaction mix is listed.

**Step 2: SNaPshot Minisequencing**

**2.1 Singleplex SNaPshot minisequencing (COMT)**

Singleplex SNaPshot minisequencing of COMT(rs4680) was performed in a total volume of 10 µL containing 3 µL of purified PCR product, 5 µL SNaPshot Multiplex Ready Reaction mix (Thermo Fisher) and 2 µL of diluted minisequencing primer (pCOMT 2 µM; details see Table 3). The cycling conditions (25 cycles) using the Veriti thermal cycler (AB) were as follows: denaturation at 96 °C for 10 seconds, annealing at 50 °C for 5 seconds and extension at 60 °C for 30 seconds.

ExoSAP-IT treatment was again applied for the clean-up of the minisequencing reaction. 5 µl of purified reaction product was then mixed with 9.3 µL Hi-Di formamide (AB) and 0.2 µL of GeneScan-LIZ 120 internal size standard (AB). After a denaturing step for 5 min at 98 °C followed by cooling to 4 °C the fragments were separated on an ABI PRISM 310 Genetic Analyzer (AB) with POP4 polymer and analysed with GeneMapper v3.2 software. Calling of SNP variants based on minisequencing was in full agreement to results from direct sequencing of PCR products in selected DNA samples.

**2.2 Multiplex SNaPshot (ANKK1, BDNF, CDH13, OPRM1, PPP1R1B)**

Multiplex SNaPshot minisequencing was performed in a total volume of 10 µL containing 3 µL of purified PCR product, 5 µL SNaPshot Multiplex Ready Reaction mix (Thermo Fisher) and 2 µL multiplex primers mix (1 µM of pBDN; 2 µM of pANKK1, pCDH13, pOPRM1 and PPP1R1B; details provided in Table 3). Thermal cycling conditions (28 cycles) were as follows: denaturation at 96 °C for 10 seconds, annealing at 58 °C for 5 seconds and extension at 60 °C for 30 seconds. ExoSAP-IT clean-up and automated detection of minisequencing products by capillary electrophoresis was conducted using the same conditions as for the singleplex SNaPshot (details provided in 2.1). Calling of SNP variants based on minisequencing was in full agreement to results from direct sequencing of PCR products in selected DNA samples.

**Table 3 Minisequencing primer information**

| Primer name^a^ | SNP (Alleles)  Location^b^ | Primer sequence 5’-3’^c^ | Primer binding site^d^ |
| --- | --- | --- | --- |
| pCOMT | rs4680 (G/A)  chr22:19963748 | *(GATC)_4_*GGATGGTGGATTTCGCTGGC | chr22:19963728-19963747(+) |
| pANKK1 | rs1800497 (C/T)  chr11:113400106 | CCATCCTCAAAGTGCTGGTC | chr11:113400086-113400105(+) |
| pBDNF | rs6265 (G/A)  chr11:27658369 | *(GATC)_8_*TCATTGGCTGACACTTTCGAACAC | chr11:27658370-27658393(-) |
| pCDH13 | rs3784943 (G/A)  chr16:83678830 | *(GATC)_10_*CCTACTTTGTCATCAGCACTGCTTT | chr16:83678805-83678829(+) |
| pOPRM1 | rs1799971 (G/A)  chr6:154039662 | CGCATGGGTCGGACAGGT | chr6:154039663-154039680(-) |
| pPPP1R1B | rs907094 (C/T)  chr17:39634118 | *(GATC)_6_*GTATACTCAAGGAGGACCCACAG | chr17:39634119-39634141(-) |

^a^ COMT primer was only used in singleplex SNaPshot; all other primers were combined in a multiplex SNaPshot.

^b^ Chromosome number and genomic location of target single nucleotide polymorphism (SNP) according to UCSC version hg38 (<http://genome.ucsc.edu/>).

^c^ The non-specific primer tail is underlined in Italics

^d^ Chromosome number and genomic location of primer binding site (orientation provided in brackets) according to UCSC version hg38.
